# Promyelocytic leukemia (PML) suppresses hepatitis B virus cccDNA transcription through SUMOylation- and SMC5/6-dependent mechanisms

**DOI:** 10.64898/2026.10.05.756908

**Authors:** Jiapeng Wu, Feng Li, James Ahodantin, Masaya Funaki, Yaxu Wu, Lishan Su

**Author notes:** J.W. and F.L. contributed equally to this work. Correspondence: Lishan Su, PhD.

## Abstract

Chronic hepatitis B virus (HBV) infection persists under current antiviral therapy because covalently closed circular DNA (cccDNA) remains in the infected hepatocytes as a durable transcriptional template. Although type I interferon (IFN-I) can reduce HBV cccDNA gene expression, the host effectors and mechanisms that restrict cccDNA remain incompletely defined. Using a minicircle-based HBV cccDNA expressing a Gaussia luciferase reporter gene (mcHBV-GLuc), we investigated how promyelocytic leukemia protein (PML) suppresses HBV cccDNA activity. Multiple PML isoforms reduced cccDNA-driven reporter activity and HBsAg production, and HBx-deficient (ΔX) mcHBV cccDNA was more susceptible than wild-type cccDNA to PML-mediated inhibition. Interestingly, efficient antiviral activity required the major PML SUMOylation sites. Disruption of SMC5/6 function attenuated PML-mediated repression of HBx-deficient cccDNA, associated with reduced PML SUMOylation and PML nuclear-body organization. These findings identify PML as a SUMO-dependent inhibitor of HBV cccDNA activity and reveal a context-dependent functional relationship between PML and SMC5/6 that is modulated by HBx.

## Introduction

Chronic hepatitis B (CHB) remains a major cause of cirrhosis and hepatocellular carcinoma, with an estimated 254 million people living with chronic hepatitis B virus (HBV) infection worldwide and more than one million HBV-attributable deaths in 2022 [1]. Nucleos(t)ide analogues (NUC) efficiently inhibit the viral reverse transcriptase and can maintain profound suppression of serum HBV DNA, but they do not directly eliminate the nuclear viral transcription template and therefore usually require long-term therapy. The central therapeutic goal is a functional cure, generally defined as sustained hepatitis B surface antigen (HBsAg) loss with or without anti-HBs seroconversion after treatment withdrawal [2]. Pegylated interferon-alpha (PEG-IFN-alpha) offers a finite immunomodulatory regimen and can produce durable off-treatment control in a subset of patients, as established in pivotal HBeAg-positive and HBeAg-negative clinical trials, but its use is constrained by toxicity, contraindications, and highly variable response rates [3, 4]. These limitations have shifted the field from simple suppression of virion production toward understanding the viral and host mechanisms that maintain or silence the persistent HBV reservoir.

HBV persistence during NUC treatment is sustained principally by covalently closed circular DNA (cccDNA), a stable episomal molecule formed in the hepatocyte nucleus after repair of incoming relaxed circular DNA [5]. The precursor-product relationship between cytoplasmic relaxed circular DNA and the amplified nuclear cccDNA pool was first demonstrated in duck hepatitis B virus-infected hepatocytes [6]. Subsequent ultrastructural and biochemical work showed that nuclear HBV DNA is assembled with histones into a nucleosome-containing minichromosome [7]. cccDNA therefore behaves as a chromatinized transcriptional template whose activity is governed by histone modifications, nucleosome organization, viral proteins, and host transcriptional regulators [8, 9]. HBx is a major determinant of this transcriptional competence: HBx associates with the cccDNA minichromosome and promotes a permissive epigenetic state, whereas HBx-deficient cccDNA becomes rapidly repressed [10]. Because cccDNA can persist despite potent inhibition of reverse transcription, and because integrated HBV DNA is unable to generate infectious HBV virions, durable HBV cure will likely require both deeper silencing or elimination of cccDNA and restoration of effective antiviral immunity [2, 5, 8].

HBV has historically been described as a stealth virus because the early phase of infection elicits little detectable hepatic type I interferon (IFN-I) induction compared with many other viral infections [11, 12]. HBV replication is unusually associated with a muted or spatially restricted innate response, although cytokines produced by infected hepatocytes, Kupffer cells, natural killer cells, and virus-specific T cells can exert powerful antiviral effects once appropriately induced. Foundational studies in HBV transgenic mice showed that cytotoxic T lymphocytes can inhibit viral gene expression and replication without hepatocyte destruction, through noncytolytic mechanisms mediated in large part by IFN-gamma and tumor necrosis factor-alpha [13].

IFN-alpha, IFN-gamma, and TNF-alpha can accelerate the loss of HBV transcripts and reduce viral replication through both transcriptional and post-transcriptional pathways [14-16]. Work spanning transgenic animals and hepatocyte models has identified several mechanistically distinct points at which IFN-induced genes (ISGs) restrict HBV. The dynamin-like GTPase MxA inhibits HBV by interfering with post-transcriptional utilization and nuclear export of viral RNAs containing the post-transcriptional regulatory element [17]. TRIM22, an IFN-inducible RING-finger E3 ligase, represses the HBV core promoter in a nuclear RING-domain-dependent manner [18]. The 3’-5’ exonuclease ISG20 binds the epsilon stem-loop of HBV RNA and directly degrades viral transcripts [19]; later work showed that ISG20 preferentially recognizes N6-methyladenosine-modified HBV RNAs in cooperation with the m6A reader YTHDF2 [20]. IFI16 provides another nuclear layer of restriction by sensing episomal HBV DNA and coupling innate signaling to epigenetic repression of cccDNA [21]. Collectively, these studies show that the IFN response is distributed across multiple viral replication steps and cellular compartments, but they also underscore that the identity and organization of the nuclear ISG effectors acting directly on cccDNA remain incompletely resolved.

A particularly important advance was the demonstration that interferons act directly on the cccDNA minichromosome. Belloni showed that IFN-alpha suppresses cccDNA transcription in cell culture and human-liver chimeric mice by inducing histone hypoacetylation and recruiting repressive chromatin regulators to cccDNA [22]. Independent work in other hepadnavirus models similarly found that IFN-alpha alters epigenetic modifications on cccDNA minichromosomes and reduces transcription without requiring loss of the DNA template [23]. In addition to transcriptional silencing, certain cytokines can promote loss of cccDNA. IFN-gamma and TNF-alpha released by activated T cells can likewise induce APOBEC3-dependent cccDNA decay without direct cytolysis [24]. More recently, ISG20 was shown to cooperate functionally with APOBEC3A for efficient IFN-induced cccDNA loss, linking RNA exonuclease activity to degradation of the persistent cccDNA [25].

Promyelocytic leukemia protein (PML) functions at the intersection of interferon signaling, nuclear organization, and intrinsic antiviral defense. Two independent studies first demonstrated in 1995 that PML expression and PML nuclear bodies (PML-NBs) increase rapidly after interferon treatment [26, 27]. Direct antiviral activity was subsequently established by showing that interferon-induced PML confers resistance to RNA viruses [28], and extensive work in herpesviruses and other DNA viruses has defined PML nuclear bodies as dynamic platforms that detect, sequester, chromatinize, or silence foreign nuclear DNA [29, 30]. PML isoforms share an N-terminal RING-B-box-coiled-coil domain that supports oligomerization but contain distinct C-terminal regions that can specify partner recruitment and antiviral activity. PML nuclear-body maturation is controlled by the small ubiquitin-like modifier (SUMO) system. PML is conjugated at three major SUMO-acceptor lysines, K65, K160, and K490 [31, 32], and multivalent interactions between SUMOylated PML and SUMO-interacting motifs drive recruitment and retention of client proteins within mature nuclear bodies [33]. These properties make PML an IFN-inducible scaffold capable of converting a soluble ISG response into a spatially organized nuclear restriction compartment.

Two independent studies in 2016 identified the structural maintenance of chromosomes 5/6 (SMC5/6) complex as the host restriction factor targeted by HBx: HBx recruits DDB1-CUL4 ubiquitin-ligase machinery to degrade SMC5/6 and thereby relieve repression of episomal HBV transcription [34, 35]. Niu et al. subsequently reported that SMC5/6 restricts HBV when localized to nuclear domain 10 (ND10), associated with PML nuclear bodies, and that depletion of PML or other ND10 components alters SMC6 localization and partially restores transcription of HBx-deficient HBV [36]. More recently, SLF2 was shown to interact with SMC5/6 and direct transcriptionally inactive HBV episomal DNA to PML bodies, providing a molecular bridge between viral DNA recognition, SMC5/6 recruitment, and spatial sequestration at ND10 [37]. These studies establish PML bodies as an important site of SMC5/6-mediated restriction, but they do not resolve whether PML SUMOylation and SMC5/6 function are mechanistically interdependent.

The SMC5/6 complex also contains the conserved SUMO E3 ligase NSMCE2 (Nse2/MMS21). Nse2/MMS21 was independently identified as an integral SMC5/6-associated SUMO ligase that can promote SUMOylation of SMC5/6 components and other chromatin-associated substrates [38-40]. This enzymatic activity provides a plausible biochemical connection between SMC5/6 and the SUMO-rich PML nuclear bodies. Supporting such spatial coupling, NSE2-dependent SUMOylation of topoisomerase IIα has been observed at PML bodies [41].

We report here that multiple PML isoforms inhibit HBV cccDNA activity through HBx-related mechanisms and that efficient antiviral activity requires PML SUMOylation. Functional epistasis experiments further show that inhibition of SMC5/6 with dominant-negative SMC6 K82E [34] significantly relieves PML-mediated repression of HBx-deficient cccDNA. Finally, we show that SMC5/6 function supports PML SUMOylation and proper PML-NB organization. Together, these findings place PML, HBx and SMC5/6 in a coordinated but context-dependent SUMO-regulated antiviral pathway.

## Materials and Methods

### Cells and culture conditions

HepG2 cells were maintained in Dulbecco’s modified Eagle medium supplemented with 10% fetal bovine serum and 1% penicillin-streptomycin at 37 °C and 5% CO2. Cells were seeded to reach the appropriate confluence at the time of transfection.

### Plasmids and minicircle HBV cccDNA reporter system

The mcHBV-GLuc cccDNA reporter was generated from a parental pre-minicircle construct containing a full-length HBV genome engineered to express Gaussia luciferase through a self-cleaving 2A peptide [42]. Minicircle recombination removed bacterial backbone sequences, including the pUC origin and antibiotic-resistance cassette, to generate an approximately 4-kb circular HBV reporter. WT and HBx-deficient (ΔX) mcHBV-GLuc cccDNA constructs were used to assess HBx-dependent regulation. Expression plasmids encoded the indicated ISGs, PML-3, PML-4, PML-6, FLAG-PML4, HA-PML-3, HA-PML-3 ΔSUMO (K65R/K160R/K490R), Myc-SUMO3, and V5-SMC6 K82E.

### Transfection

HepG2 cells were transfected with Lipofectamine 3000 according to the manufacturer’s instructions. Unless otherwise indicated, 1 μg total DNA was used per well containing approximately 1 × 10^6 cells. Empty-vector DNA was added as needed to keep the total amount of transfected DNA constant across conditions.

### Gaussia luciferase assays

Secreted Gaussia luciferase activity was measured in culture supernatants using the Pierce Gaussia Luciferase Glow Assay Kit (catalog no. 16160) according to the manufacturer’s instructions. For most assays, 50 μl of supernatant was analyzed. Relative light units were recorded with a plate luminometer and normalized to the appropriate mock, empty-vector, or matched control condition. Percentage inhibition was calculated relative to the corresponding control within each experiment.

### HBsAg ELISA

HBsAg in culture supernatants was measured using a quantitative hepatitis B surface antigen ELISA kit (catalog no. 4110) according to the manufacturer’s protocol. Supernatants were collected at the indicated time points, including 72 hours post-transfection for the PML isoform and SUMOylation experiments.

### SMC6 K82E overexpression experiments

For SMC5/6 dependency assays, HepG2 cells were first transfected with WT or ΔX mcHBV-GLuc cccDNA. Three days later, cells were transferred to 96-well plates and transfected with empty vector, V5-SMC6 K82E [34], PML-4, or the combination of SMC6 K82E and PML-4. GLuc activity was measured at 24, 48, and 72 hours after the SMC/PML transfection. SMC6 K82E was also tested alone at 72 hours to determine its effect on basal host restriction in the absence of ectopic PML.

### Immunofluorescence microscopy and PML-NB morphology scoring

To assess PML nuclear-body organization, HepG2 cells were transfected with FLAG-PML4 alone or co-transfected with FLAG-PML4 and V5-SMC6 K82E and processed for immunofluorescence imaging. FLAG-PML was visualized in red, V5-SMC6 K82E in green, and nuclei were counterstained with DAPI (blue). Images were acquired at 20× magnification. Four non-overlapping PML-only fields and four non-overlapping co-transfection fields were analyzed. PML-positive cells were classified as punctate PML-NB-positive when discrete PML puncta were associated with a DAPI-stained nucleus, or as abnormal/ring-like when PML formed a broad nuclear/perinuclear ring, diffuse non-punctate signal, or otherwise lacked discrete PML-NB morphology. In co-transfected cultures, PML-positive cells were further stratified as SMC6-K82E-positive or SMC6-K82E-negative according to the presence or absence of detectable green signal in the same cell. Cells truncated by the field boundary or with ambiguous signal were excluded.

### Immunoblotting and immunoprecipitation

Cells were lysed in RIPA buffer supplemented with protease and phosphatase inhibitor cocktails and protein concentrations were determined by BCA assay. Whole-cell lysates were resolved by 4-12% SDS-PAGE and transferred to nitrocellulose membranes. Blots were probed with antibodies against FLAG (Sigma-Aldrich, F1804-1MG), HA (Sigma-Aldrich, SAB2702196), Myc (ProteinTech, 16286-1-AP), SMC5 (ProteinTech, 14178-1-AP), SMC6 (ProteinTech, 14465-1-AP), HBc (Dako, B0586), and β-actin. Horseradish peroxidase-conjugated secondary antibodies were used for detection, and β-actin served as a loading control. Signals were visualized by enhanced chemiluminescence using a Bio-Rad CCD imaging system and quantified with ImageJ. For PML SUMOylation assays, FLAG-PML4 (Sigma-Aldrich, FLAGIPT1-1KT) or HA-PML-3 (Sigma-Aldrich, IP0010-1KT) was immunoprecipitated from clarified lysates, and immunoprecipitates were analyzed by tag-specific immunoblotting. Corresponding whole-cell inputs were analyzed in parallel.

### Statistical analysis

Data are presented as mean ± SEM unless otherwise indicated. Multiple-group comparisons were performed by one-way ANOVA followed by Fisher’s LSD post-hoc test using Prism 10 (GraphPad Software). For the PML-NB morphology analysis, each plotted point represents the percentage of classifiable PML-positive cells with punctate PML-NB morphology in one imaging field (n = 4 fields per group); pooled cell counts were reported separately and were not treated as independent observations for significance testing. Statistical significance is indicated as follows: *P < 0.05; **P < 0.01; ***P < 0.001; ****P < 0.0001; ns, not significant.

## Results

### Multiple PML Isoforms Suppress HBV cccDNA Activity, with Greater Restriction in the Absence of HBx

To investigate host factors that regulate HBV cccDNA transcription, we used a minicircle HBV reporter (mcHBV-GLuc cccDNA) engineered to express Gaussia luciferase (GLuc) through a self-cleaving 2A peptide (Fig. S1) [42]. The resulting minicircle supports transcription of pregenomic and subgenomic RNAs, while secreted GLuc provides a quantitative surrogate for cccDNA transcriptional output.

We investigated how PML nuclear-body proteins inhibit cccDNA-driven transcription using the mcHBV-GLuc cccDNA model. We confirmed the inhibition with three nuclear PML isoforms, PML-3, PML-4, and PML-6, whose exon structures are summarized in Fig. S2A. HepG2 cells were co-transfected with mcHBV-GLuc cccDNA and the indicated PML expression plasmids. Each isoform significantly reduced GLuc activity at 24, 48, and 72 hours post-transfection relative to the vector control (Fig. 1A-C), with the largest absolute differences observed at the later time points as reporter activity accumulated. Consistent with suppression of cccDNA-driven gene expression, all three PML isoforms also markedly reduced HBsAg secretion at 72 hours post-transfection (Fig. 1D). These data show that antiviral activity is shared by multiple tested nuclear PML isoforms.

**Figure 1.**
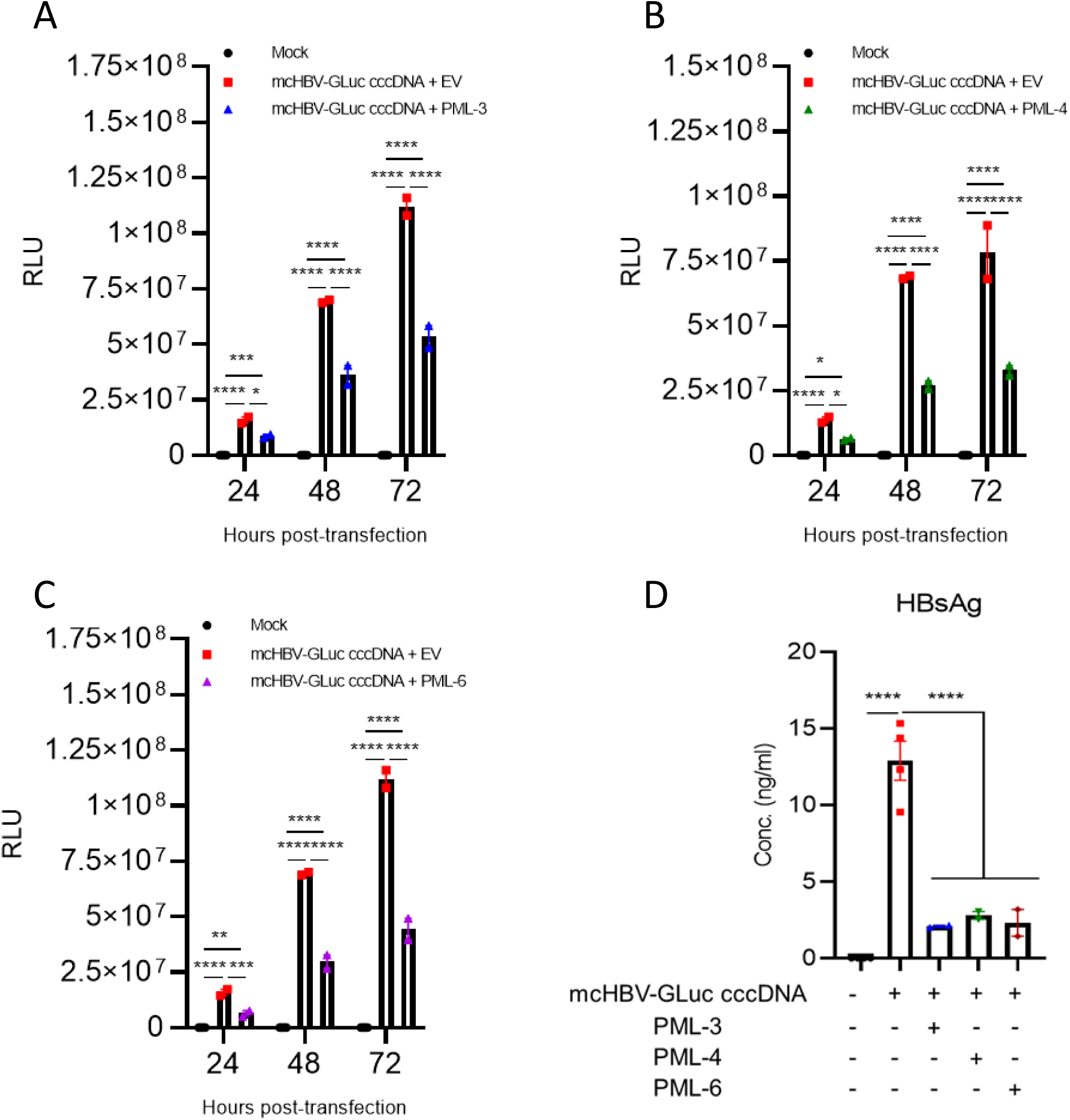
PML isoforms suppress HBV cccDNA activity. (A-C) HepG2 cells were co-transfected with mcHBV-GLuc cccDNA and empty vector (EV), PML-3 (A), PML-4 (B), or PML-6 (C). GLuc activity in culture supernatants was measured at 24, 48, and 72 hours post-transfection and is presented as relative light units (RLU). Mock-transfected cells are shown as a background control. (D) HBsAg concentrations in culture supernatants were measured by ELISA at 72 hours post-transfection. Data are shown as mean ± SEM. Statistical analysis was performed by one-way ANOVA followed by Fisher’s LSD post-hoc test (*P < 0.05; **P < 0.01; ***P < 0.001; ****P < 0.0001).

To determine whether HBx modulates PML restriction, HepG2 cells were co-transfected with wild-type (WT) or HBx-deficient (ΔX) mcHBV-GLuc cccDNA together with empty vector, PML-3, or PML-4. PML-3 and PML-4 reduced reporter activity from both WT and ΔX cccDNA, but ΔX cccDNA displayed significantly greater percentage inhibition by either PML isoform (Fig. 2A-B). A 24- to 72-hour time course further showed increasing separation between the empty-vector and PML-4 conditions for ΔX cccDNA, consistent with progressive PML-mediated restriction in the absence of HBx (Fig. 2C). Thus, HBx partly counteracted the antiviral activity of PML.

**Figure 2.**
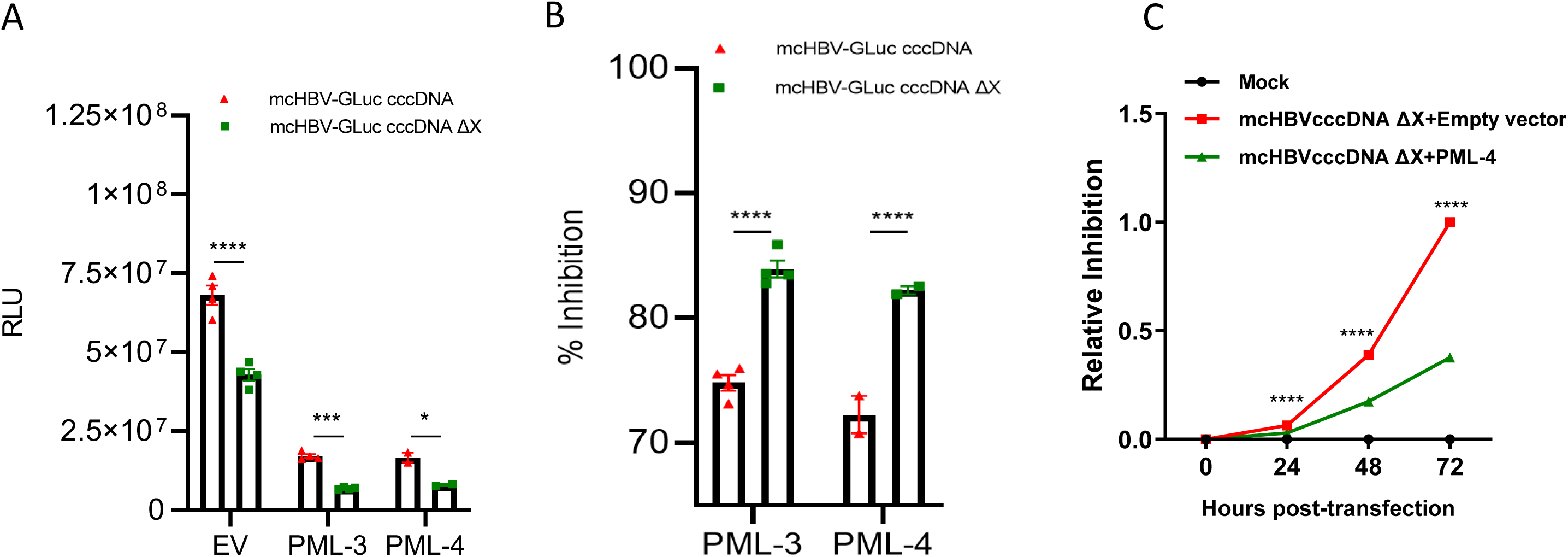
PML inhibits HBV cccDNA through HBx-related mechanisms. (A) HepG2 cells were co-transfected with WT or HBx-deficient (ΔX) mcHBV-GLuc cccDNA and EV, PML-3, or PML-4. GLuc activity was measured at 72 hours post-transfection and is presented as RLU. (B) Percentage inhibition mediated by PML-3 or PML-4 was calculated relative to the matched EV control for WT and ΔX cccDNA. (C) Time-course analysis of ΔX mcHBV-GLuc cccDNA in cells transfected with EV or PML-4. Reporter activity was assessed at 0, 24, 48, and 72 hours and expressed relative to the matched control. Data are shown as mean ± SEM. Statistical analysis was performed by one-way ANOVA followed by Fisher’s LSD post-hoc test (*P < 0.05; ***P < 0.001; ****P < 0.0001).

### PML SUMOylation Is Required for the Suppression of HBV cccDNA

PML nuclear-body assembly and antiviral function are dependent on SUMOylation. To test whether PML SUMOylation is required for HBV cccDNA repression, we constructed a SUMOylation-defective PML-3 mutant in which the three major SUMO-acceptor lysines were replaced by arginine (PML-3 ΔSUMO; K65R/K160R/K490R; Fig. S2B). Wild-type PML-3 strongly reduced mcHBV-GLuc activity, whereas PML-3 ΔSUMO showed little or no antiviral activity relative to the control vector condition (Fig. 3A-B). Input immunoblotting confirmed equal expression of both PML proteins and Myc-SUMO3 (Fig. 3C), while anti-HA immunoprecipitation demonstrated abundant HA-tagged SUMO-conjugated species for wild-type PML-3 but abolished SUMOylation of PML-3 ΔSUMO (Fig. 3D). Consistent with the gLuc reporter expression, wild-type PML-3 strongly reduced HBsAg secretion, whereas the ΔSUMO mutant had no significant effect (Fig. 3E). These results demonstrate that PML SUMOylation is required for efficient suppression of HBV cccDNA activity.

**Figure 3.**
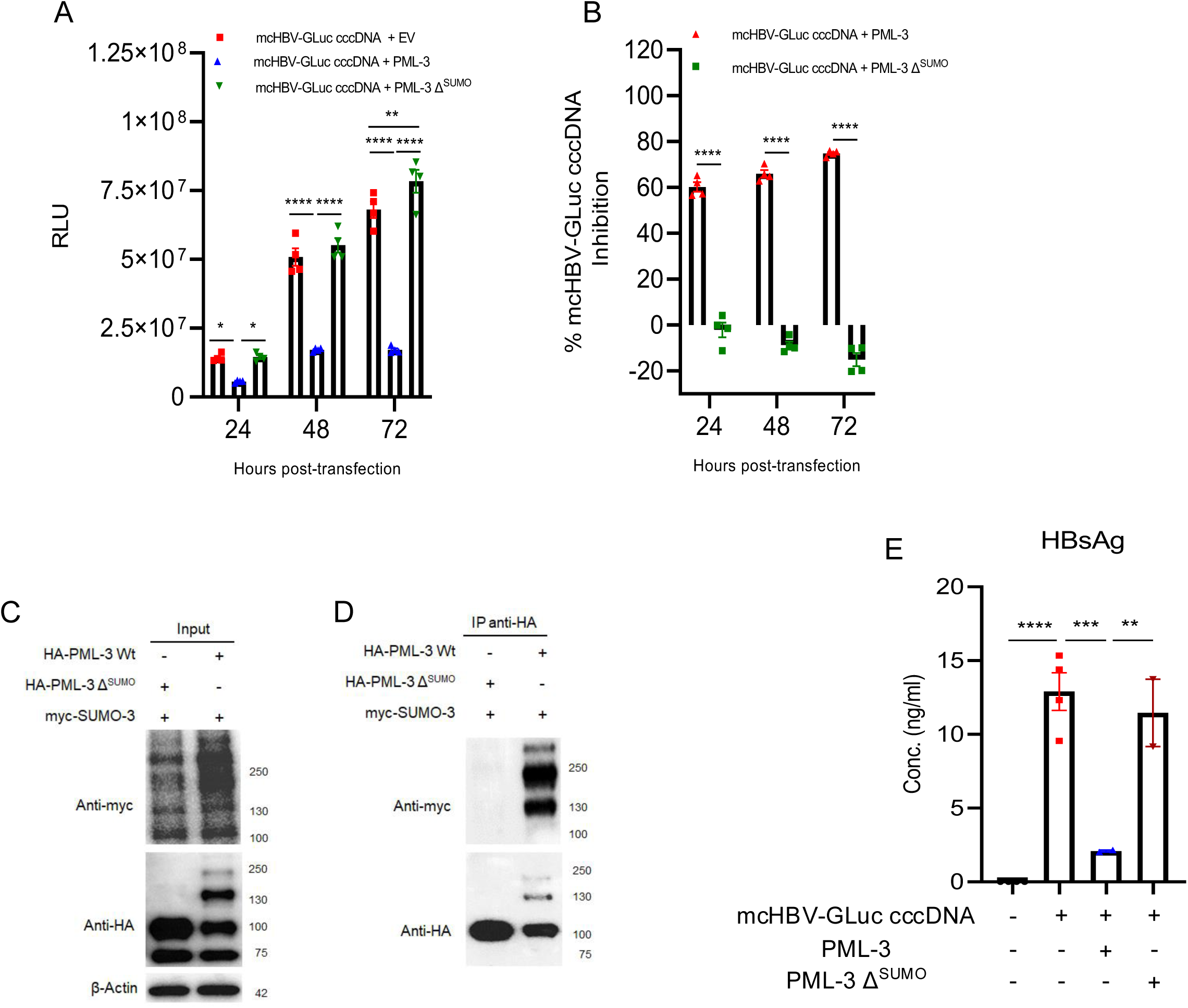
PML suppression of HBV cccDNA requires SUMOylation. HepG2 cells were co-transfected with mcHBV-GLuc cccDNA and EV, wild-type PML-3, or SUMOylation-defective PML-3 ΔSUMO (K65R/K160R/K490R). (A) GLuc activity was measured at 24, 48, and 72 hours post-transfection and is presented as RLU. (B) Percentage inhibition by PML-3 or PML-3 ΔSUMO was calculated relative to the matched EV control at each time point. (C) Input immunoblot confirming expression of HA-PML-3, HA-PML-3 ΔSUMO, and Myc-SUMO3. (D) Anti-HA immunoprecipitation followed by anti-Myc and anti-HA immunoblotting to detect SUMO3-conjugated PML. (E) HBsAg concentrations in culture supernatants at 72 hours post-transfection. Data are shown as mean ± SEM. Statistical analysis was performed by one-way ANOVA followed by Fisher’s LSD post-hoc test (*P < 0.05; **P < 0.01; ***P < 0.001; ****P < 0.0001).

### SMC5/6 Is Required for PML-Mediated Repression of HBx-Deficient cccDNA

To determine whether PML-mediated repression of HBx-deficient cccDNA depends on functional SMC5/6, HepG2 cells were first transfected with ΔX mcHBV-GLuc cccDNA. Three days later, cells were transferred to 96-well plates and transfected with control vector, PML-4 with and without the dominant-negative SMC6 K82E [34]. GLuc activity was measured at 24, 48, and 72 hours after the SMC/PML transfection (Fig. 4A). At 72 hours, while SMC6 K82E enhanced ΔX cccDNA activity by inhibiting SMC5/6 activity as we have reported [34], the ability of PML-4 to suppress ΔX cccDNA was significantly counteracted by SMC6 K82E (Fig. 4B). Earlier measurements at 24 and 48 hpt showed the same directional effect, with a less pronounced rescue than at 72 hpt (Fig. S3B-C). The data show that disruption of SMC5/6 function counteracts PML-mediated repression of ΔX cccDNA, supporting a requirement for functional SMC5/6. In the presence of HBx, with SMC5/6 degraded, SMC6 K82E showed no effect on the PML activity (Fig. S3D-F). SMC6 K82E alone did not derepress WT cccDNA. In contrast to ΔX cccDNA, PML-4 suppressed WT cccDNA activity even when SMC6 K82E was co-expressed.

**Figure 4.**
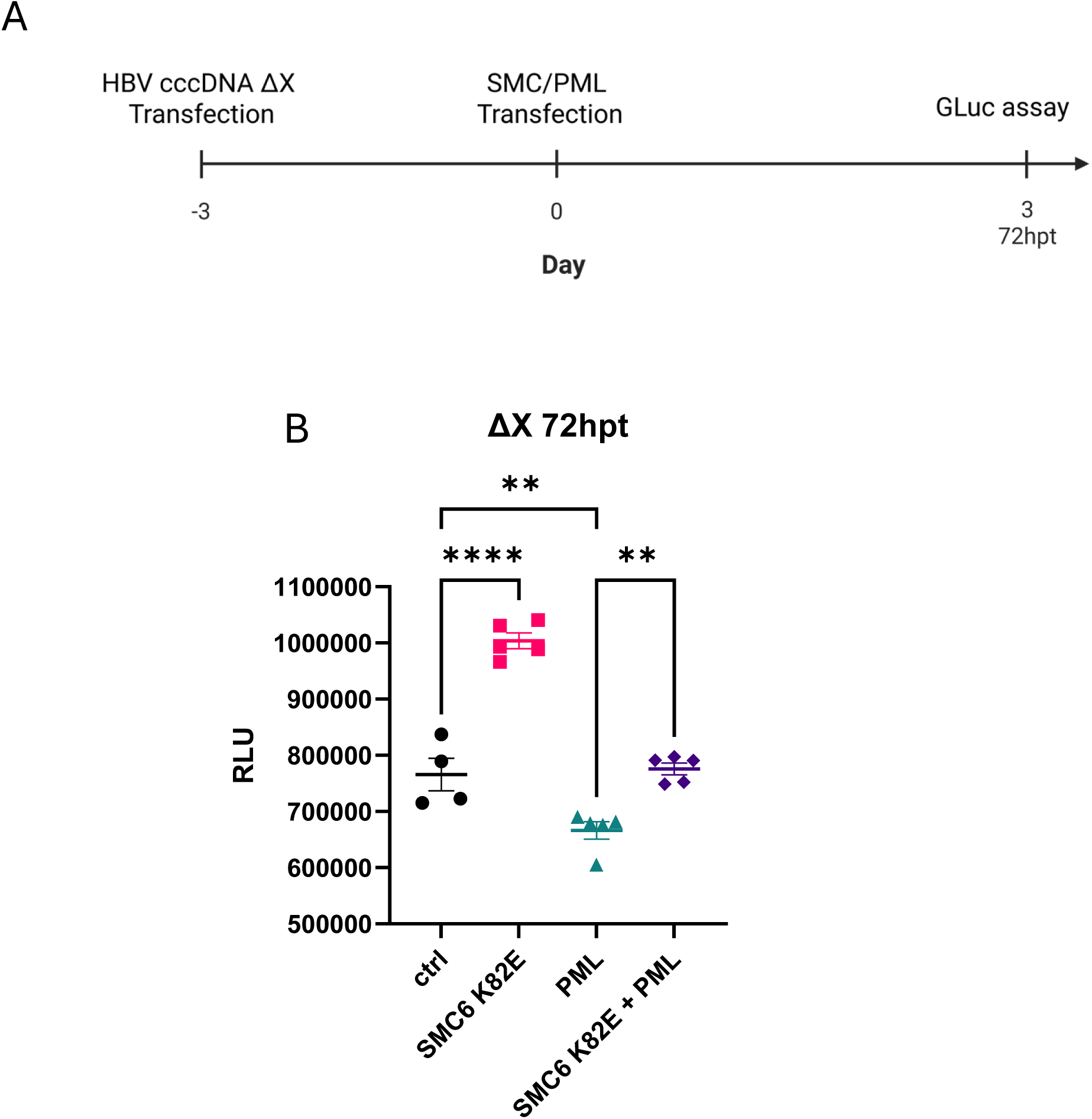
SMC5/6 is required for sustained PML-mediated repression of HBx-deficient cccDNA. (A) Experimental timeline. HepG2 cells were transfected with HBx-deficient (ΔX) mcHBV-GLuc cccDNA on day -3. On day 0, cells were transferred to 96-well plates and transfected with control vector, SMC6 K82E, PML-4, or SMC6 K82E plus PML-4. GLuc activity was measured 72 hours after SMC/PML transfection. (B) ΔX mcHBV-GLuc cccDNA activity at 72 hpt. Data are shown as mean ± SEM. Statistical analysis was performed by one-way ANOVA followed by Fisher’s LSD post-hoc test (*P < 0.05; **P < 0.01; ***P < 0.001; ****P < 0.0001; ns, not significant).

### SMC5/6 Supports Efficient PML SUMOylation

Because PML-mediated cccDNA repression required PML SUMOylation and the ΔX cccDNA experiments identified a requirement for functional SMC5/6, we tested whether SMC5/6 influences PML SUMOylation [38-40]. HepG2 cells were transfected with HBV WT or HBV ΔX cccDNA together with FLAG-PML4 and Myc-SUMO3, in the presence or absence of V5-SMC6 K82E. FLAG-PML4 was immunoprecipitated and SUMO3-conjugated PML species were detected by anti-Myc/SUMO immunoblotting (Fig. 5A). PML4 SUMOylation was approximately fourfold higher under ΔX than WT conditions (Fig. 5B). SMC6 K82E had little effect on PML4 SUMOylation in WT HBV-expressing cells (Fig. 5C), in which endogenous SMC5/6 is already degraded by HBx. This preservation of PML4 SUMOylation parallels the repression of WT cccDNA by PML in the presence of SMC6/K82E (Fig. S3). In contrast, SMC6/K82E substantially reduced SUMO3-conjugated PML4 with ΔX cccDNA (Fig. 5D), where endogenous SMC5/6 is not degraded. Whole-cell input blots confirmed equal expression of Myc-SUMO3 and FLAG-PML4 across the corresponding conditions (Fig. S4). Together, these biochemical and functional data support a context-dependent model in which SMC5/6 promotes PML SUMOylation and antiviral activity in the absence of HBx, although PML SUMOylation and repression can be maintained through an additional SMC5/6-independent mechanism.

**Figure 5.**
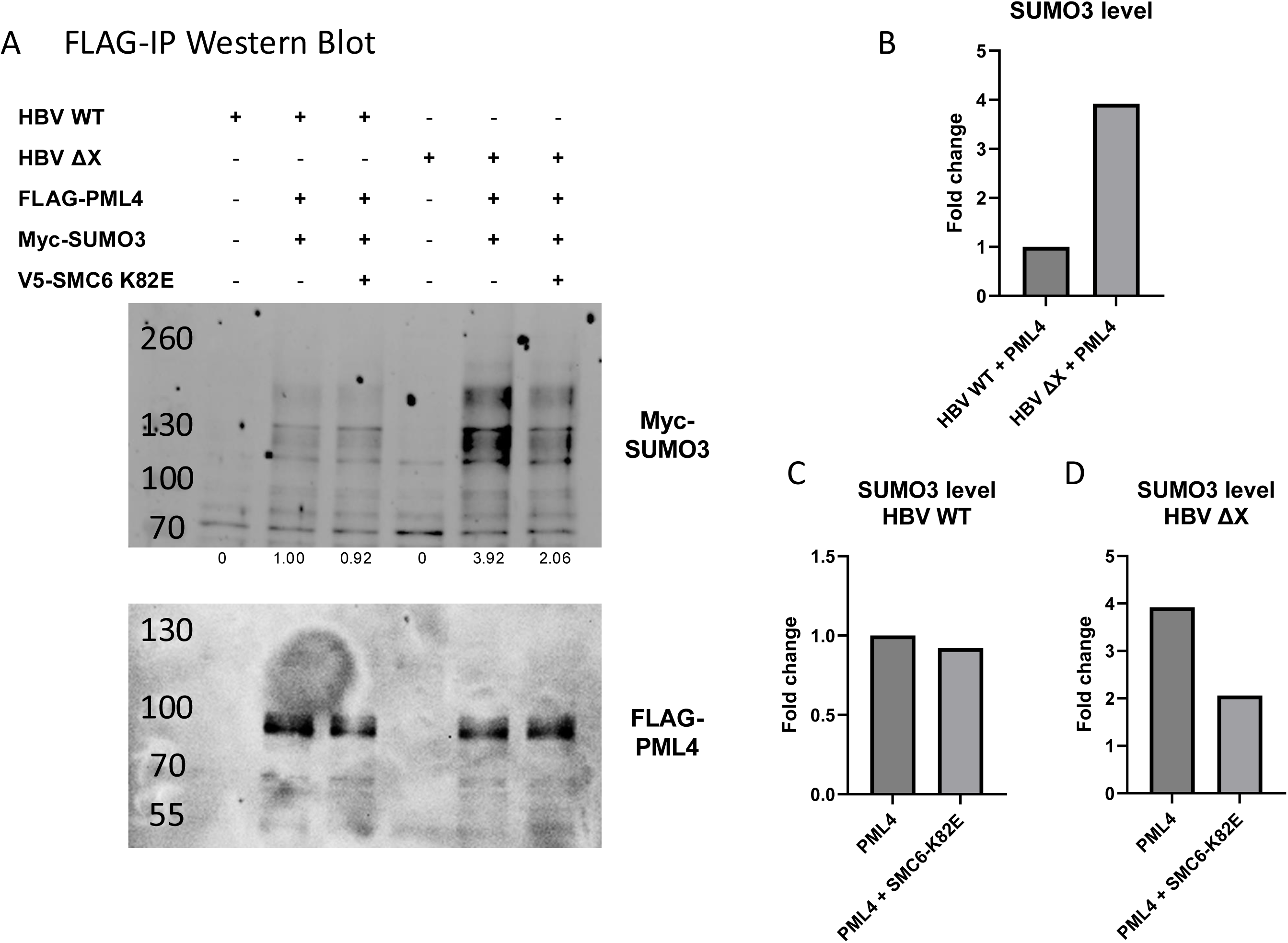
SMC5/6 promotes PML SUMOylation. HepG2 cells were transfected with HBV WT or HBV ΔX together with FLAG-PML4 and Myc-SUMO3, in the absence or presence of V5-SMC6 K82E. FLAG-PML4 was immunoprecipitated and analyzed by immunoblotting. (A) FLAG-immunoprecipitation immunoblot showing Myc-SUMO3-conjugated species and immunoprecipitated FLAG-PML4. Values below the anti-Myc blot indicate relative SUMO3 signal. (B) Relative PML4 SUMOylation under HBV WT and HBV ΔX conditions. (C) Effect of SMC6 K82E on PML4 SUMOylation in HBV WT-expressing cells. (D) Effect of SMC6 K82E on PML4 SUMOylation in HBV ΔX- expressing cells. Corresponding whole-cell input blots are shown in Fig. S4.

### Dominant-Negative SMC6 K82E Disrupts Punctate PML Nuclear-Body Organization

Because SMC5/6-dependent cccDNA restriction is linked to PML nuclear bodies, we next asked whether disruption of SMC6 function alters PML-NB organization. HepG2 cells expressing FLAG-PML4 alone showed discrete punctate PML structures associated with DAPI-positive nuclei (Fig. 6A-C). Across four analyzed fields, 63 of 99 PML-positive cells (63.6%) were classified as having punctate PML-NB morphology, whereas 36 of 99 cells (36.4%) displayed abnormal, ring-like, or otherwise non-punctate PML organization (Fig. 6K; Fig. S5A-M).

**Figure 6.**
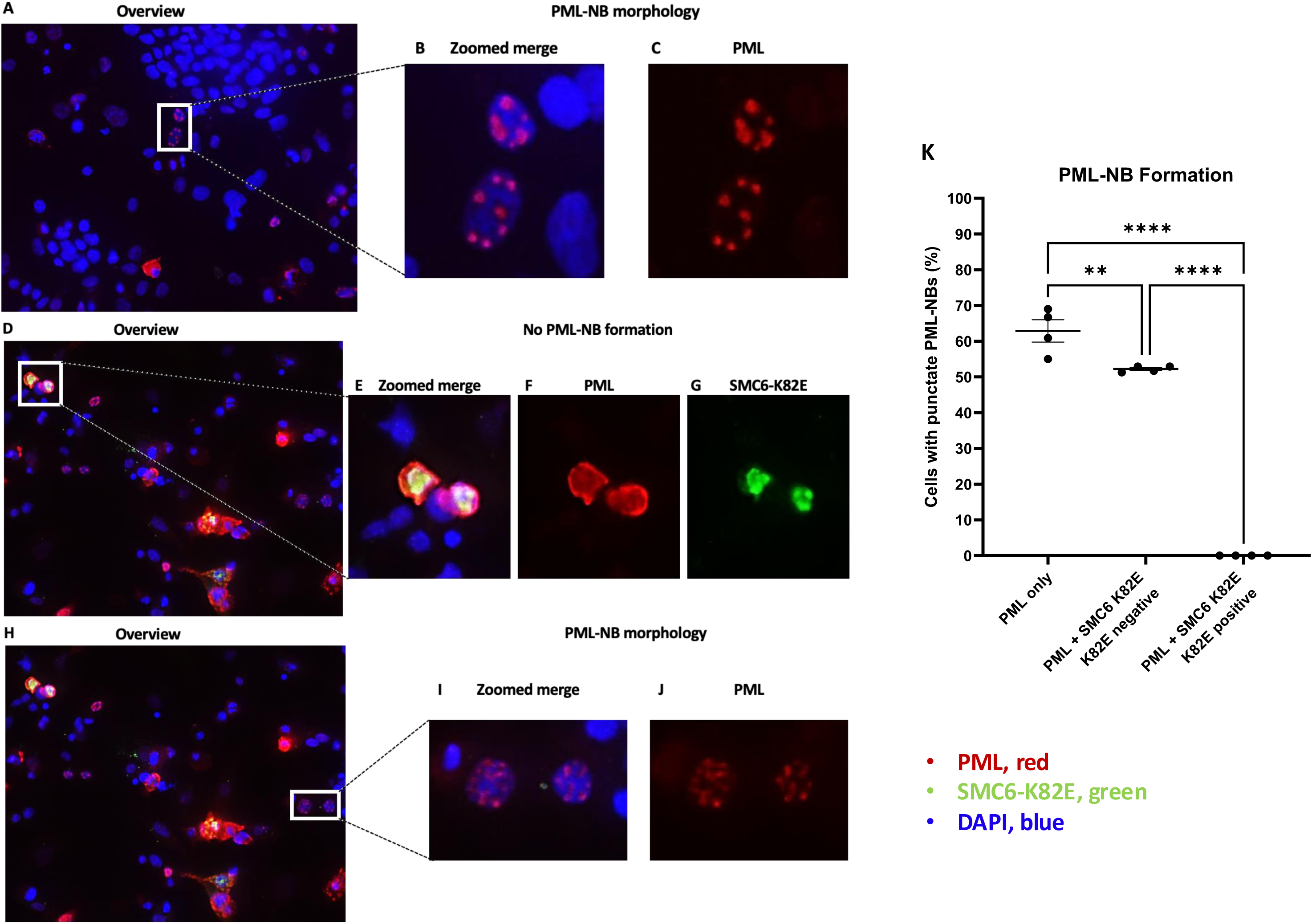
Dominant-negative SMC6 K82E disrupts punctate PML nuclear-body organization. (A) Representative merged immunofluorescence field of HepG2 cells transfected with FLAG-PML4 alone. PML is shown in red and nuclei are stained with DAPI (blue); the boxed region is enlarged in B and C. (B) Enlarged merged image and (C) corresponding PML channel showing discrete punctate PML-NB morphology. (D) Representative merged field from cells co-transfected with FLAG-PML4 and V5-SMC6 K82E; boxed PML/SMC6-K82E double-positive cells are enlarged in E-G. (E) Enlarged merge, (F) PML channel, and (G) SMC6-K82E channel showing abnormal ring-like/non-punctate PML organization in double-positive cells. (H) Representative field from the same co-transfection condition highlighting PML-positive cells lacking detectable SMC6 K82E. (I) Enlarged merge and (J) PML channel showing preserved punctate PML-NB morphology in SMC6-K82E-negative cells. (K) Field-level percentage of classifiable PML-positive cells with punctate PML-NB morphology. Each point represents one imaging field (n = 4 fields per group); bars show mean ± SEM. Statistical analysis was performed by one-way ANOVA followed by Fisher’s LSD post-hoc test (**P < 0.01; ****P < 0.0001). Images were acquired at 20× magnification; overview scale bars, 125 μm.

In cultures co-transfected with FLAG-PML4 and V5-SMC6 K82E, PML organization was strongly associated with the presence of detectable SMC6 K82E in the same cell. PML/SMC6-K82E double-positive cells showed broad ring-like or non-punctate PML patterns rather than discrete nuclear bodies (Fig. 6D-G): none of 28 double-positive cells formed punctate PML-NBs, whereas all 28 were classified as abnormal/ring-like. By contrast, neighboring PML-positive cells in the same co-transfected cultures that lacked detectable SMC6 K82E retained punctate PML-NB morphology in 52 of 100 cells (52.0%), with 48 of 100 cells (48.0%) showing abnormal/ring-like PML (Fig. 6H-J). Field-level analysis confirmed a marked reduction in punctate PML-NB-positive cells in the SMC6-K82E-positive group relative to both comparison groups (Fig. 6K). Additional fields used for scoring are shown in Fig. S5. These imaging data provide a structural correlate for the functional SMC5/6 dependency observed with ΔX cccDNA and indicate that dominant-negative SMC6 strongly disrupts normal punctate PML-NB organization.

## Discussion

In this study, we identify PML as a host restriction factor that suppresses HBV cccDNA transcription and defined its functional relationship with SUMOylation and the SMC5/6 complex. Multiple nuclear PML isoforms inhibited cccDNA-driven GLuc activity and HBsAg secretion, indicating that antiviral activity is shared across structurally distinct PML variants. The effect was stronger against HBx-deficient cccDNA and required the major PML SUMOylation sites. Importantly, the requirement for SMC5/6 was context dependent: functional SMC5/6 was required for PML-NB formation and sustained PML-mediated repression of ΔX cccDNA. Together, these findings place PML and SMC5/6 within an ND10-associated, SUMO-regulated antiviral network whose functional organization is shaped by HBx.

The greater susceptibility of ΔX cccDNA to PML-mediated inhibition is consistent with the established role of HBx as a countermeasure against SMC5/6-dependent silencing. WT cccDNA retained partial sensitivity to PML, whereas ΔX cccDNA showed both lower basal activity and greater percentage inhibition following PML expression. This pattern is consistent with HBx weakening a PML-SMC5/6 restriction pathway that is particularly effective in the absence of HBx.

The dominant-negative SMC6 K82E experiments provide functional evidence that SMC5/6 is required for PML-mediated restriction of HBx-deficient cccDNA. At 72 hpt, K82E substantially relieved PML-4-mediated repression of ΔX cccDNA (Fig. 4B), whereas the same mutant failed to restore WT reporter activity at 24, 48, or 72 hpt (Fig. S3D-F). Two non-mutually exclusive explanations could account for this difference. First, PML-4 SUMOylation remained detectable in WT cells despite SMC6 K82E (Fig. 5C), while mutation of the major PML SUMO-acceptor sites eliminated antiviral activity (Fig. 3). This raises the possibility that, when HBx is expressed, PML can remain SUMOylated and functionally active through an SMC5/6-independent SUMO-regulatory pathway. Second, HBx may reduce the effective abundance of the ectopically expressed SMC6 K82E mutant itself. HBx targets the SMC5/6 complex for CRL4-DDB1-dependent ubiquitination and proteasomal degradation [34, 35], whereas K82E is an ATPase-site mutation that disrupts SMC6 DNA binding and complex function rather than a degradation-resistant form of SMC6 [34]. Taken together, these observations raise the possibility that, when HBx is expressed, PML can remain partially SUMOylated and functionally active through an SMC5/6-independent pathway. The WT data therefore suggest that SMC5/6 is a major facilitator, but not the only pathway capable of supporting SUMO-dependent PML antiviral activity. The imaging analysis provides a structural correlate to the functional epistasis. SMC6 K82E expression was associated with a loss of punctate PML nuclear bodies, and the within-culture comparison argues that this defect tracks with dominant-negative SMC6 expression rather than the co-transfection condition itself. Although ectopic PML showed some baseline morphological heterogeneity, the overall shift in PML-NB architecture is consistent with the reduced PML4 SUMOylation observed when SMC5/6 function is disrupted.

Our immunoprecipitation results provide a biochemical correlate for this context-dependent model. PML4 SUMOylation was substantially higher under ΔX than WT conditions, and SMC6/K82E selectively reduced SUMO-conjugated PML4 in the ΔX background. These data suggest that functional SMC5/6 promotes the formation or stability of SUMOylated PML assemblies when HBx is absent. By contrast, the residual PML SUMOylation observed in WT cells was not further reduced by SMC6 K82E, consistent with the persistence of PML-mediated repression in Fig. S3. The biochemical and functional data converge on a model in which SMC5/6 is a major driver of PML SUMOylation and repression in the ΔX background, while an additional SMC5/6-independent mechanism can maintain PML SUMOylation and antiviral activity in WT HBV-expressing cells.

A potential mechanistic link between SMC5/6 and PML SUMOylation in the HBx-deficient setting is NSMCE2/Nse2 (MMS21), the SUMO E3 ligase subunit of the SMC5/6 complex. The presence of an intrinsic SUMO ligase within SMC5/6 provides a plausible mechanism through which the complex could influence the SUMO-rich environment of PML nuclear bodies [38-40]. However, the present experiments do not establish whether NSMCE2 directly modifies PML, and SMC6/K82E disrupts SMC5/6 function rather than selectively inhibiting NSMCE2 catalytic activity. Future experiments comparing wild-type and SUMO-ligase-defective NSMCE2 will therefore be required to determine whether SMC5/6 promotes PML SUMOylation through NSMCE2-dependent SUMO transfer or indirectly through preservation of PML nuclear-body organization.

These findings are consistent with prior work showing that SMC5/6 restriction depends on localization to PML nuclear bodies and that SLF2 recruits the complex and transcriptionally inactive HBV cccDNA to ND10 [36, 37]. Rather than positioning PML and SMC5/6 in a simple linear pathway, our data support a context-dependent cooperative model. In the absence of HBx, SMC5/6 provides direct episomal silencing and promotes efficient PML SUMOylation, potentially through its NSMCE2-containing SUMO machinery, thereby reinforcing a SUMO-dependent PML nuclear scaffold. When HBx depletes SMC5/6, this arm of restriction is weakened, but PML can retain residual SUMOylation and antiviral activity through an SMC5/6-independent route. HBx therefore disrupts an important component of the PML-SMC5/6 restriction network without necessarily eliminating all SUMO-dependent PML activity.

Several limitations should be considered. First, although PML is well established as an IFN-inducible protein, the present experiments did not directly test IFN-dependent induction of endogenous PML or establish that PML mediates IFN-dependent repression of HBV cccDNA. Second, PML was examined primarily by ectopic expression and a SUMOylation-defective mutant; loss-of-function studies were not performed in the mcHBV-GLuc system, and the contribution of endogenous PML to basal cccDNA repression therefore remains to be established. Third, the mcHBV-GLuc reporter is introduced by transfection, bypassing viral entry and rcDNA-to-cccDNA biogenesis, and its copy number and chromatin state may differ from infection-derived cccDNA. In addition, SMC6 K82E is a dominant-negative perturbation and may not fully phenocopy genetic depletion of the SMC5/6 complex. Finally, we did not directly measure PML or SMC5/6 occupancy on the viral episome; thus, the proposed PML-SMC5/6 relationship to cccDNA is based on functional epistasis, SUMOylation, and nuclear-body morphology rather than demonstration of a physical complex. Validation in infection-competent models using PML loss-of-function, IFN stimulation, genetic SMC5/6 perturbation, and cccDNA occupancy/localization assays will be important next steps.

In summary, PML restricts HBV cccDNA transcription through SUMO-dependent mechanisms whose reliance on SMC5/6 is shaped by HBx. In HBx-deficient cells, functional SMC5/6 is required for sustained PML-mediated repression, supports efficient PML SUMOylation, and is associated with maintenance of punctate PML-NB organization. Dominant-negative SMC6 disrupts each of these linked phenotypes: it relieves ΔX cccDNA repression, reduces PML SUMOylation, and converts PML from a punctate nuclear-body pattern to abnormal organization in double-positive cells. In WT HBV-expressing cells, where HBx reduces functional SMC5/6, PML-mediated repression and residual PML SUMOylation persist despite SMC6 K82E, suggesting an additional SMC5/6-independent route of PML activation or reduced effective K82E activity in the presence of HBx. These findings identify the PML-SMC5/6 axis as an HBx-sensitive node of intrinsic nuclear immunity and connect SMC5/6 function to both the biochemical modification and higher-order organization of PML.

## Data Availability Statement

All data generated or analyzed during this study are included in the article and its Supplementary Material, or are available from the corresponding author upon reasonable request.

## Ethics Statement

This study used the commercially available HepG2 cell line and did not involve human participants, human samples, or animals.

## Author Contributions

Conceptualization: FL and LS. Methodology: JW, FL, JA and LS. Investigation: JW, FL, JA, MF and YW. Visualization: JW, JA and LS. Resources: LS. Supervision: LS. Writing—original draft: JW. Writing—review and editing: LS.

## Funding

This work was supported by the National Institutes of Health (NIH) grant AI181664 to L.S.

## Acknowledgments

The authors thank members of the Su laboratory for helpful discussions, technical assistance, and critical feedback throughout the course of this study.

## Conflict of Interest

The authors declare that they have no competing interests.

## Supplementary Material

**Figure S1.**
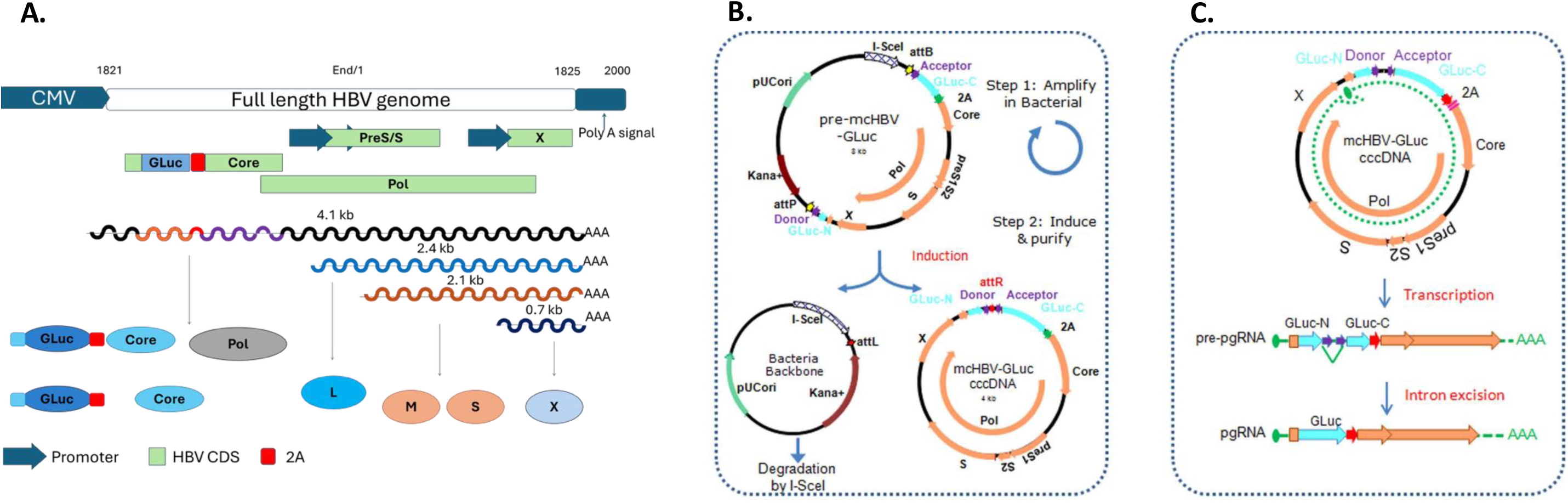
A functional mcHBV-GLuc cccDNA model. (A) Schematic of the full-length HBV genome engineered to express GLuc through a self-cleaving 2A peptide. (B) Minicircle production strategy. Site-specific recombination removes the bacterial backbone, including the pUC origin and kanamycin-resistance cassette, to generate an approximately 4-kb circular HBV reporter. (C) Functional model in which the circular mcHBV-GLuc cccDNA serves as a template for pregenomic and subgenomic RNAs, enabling quantitative assessment of cccDNA transcription through secreted GLuc activity.

**Figure S2.**
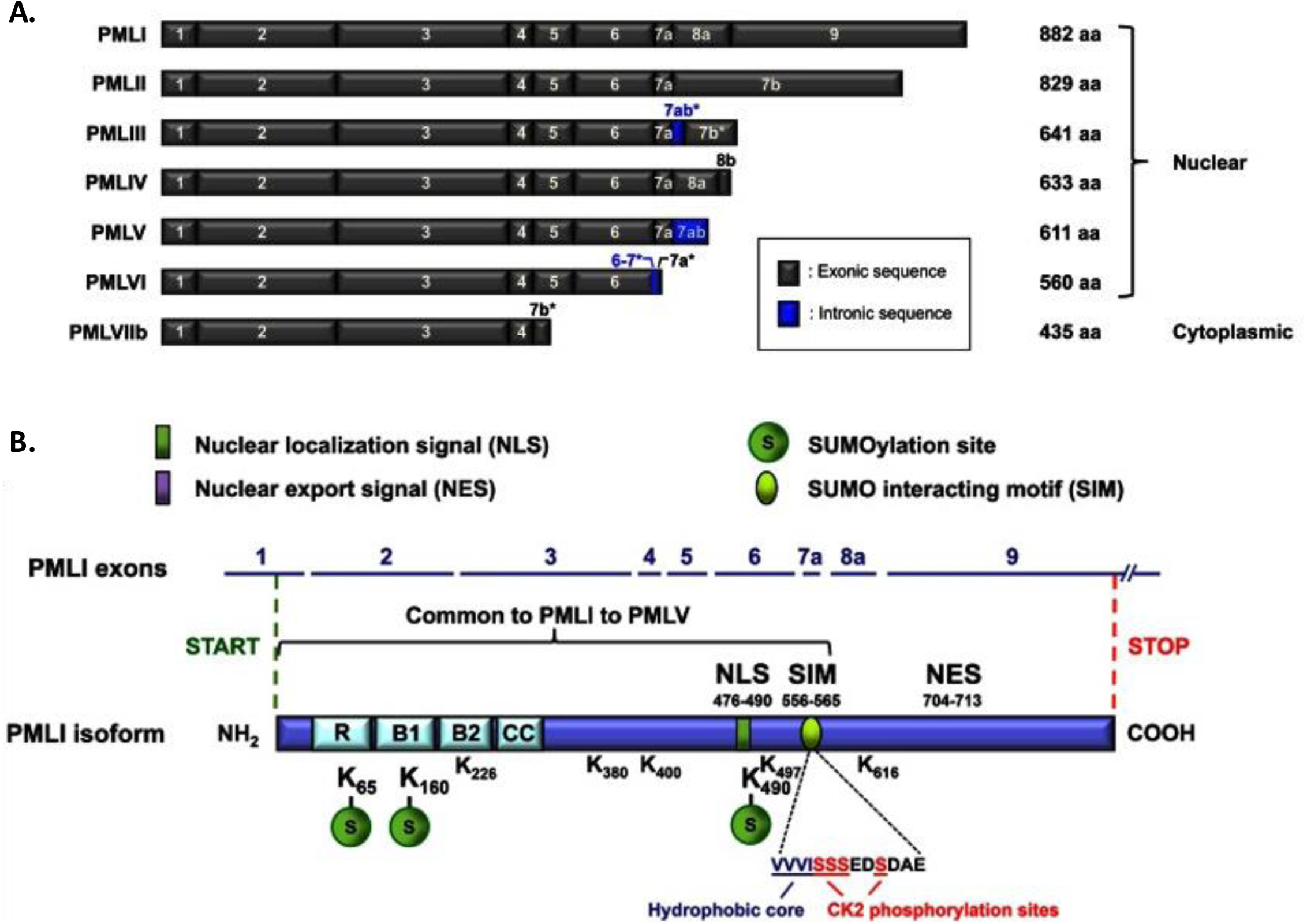
PML isoform architecture and design of the SUMOylation-defective PML mutant. (A) Schematic comparison of the major PML isoforms. Red arrows indicate the PML-3, PML-4, and PML-6 isoforms examined in this study. (B) Domain organization of PML and positions of the three major SUMO-acceptor residues, K65, K160, and K490 [32]. These lysines were mutated to arginine to generate PML-3 ΔSUMO (K65R/K160R/K490R). NLS, nuclear localization signal; SIM, SUMO-interacting motif.

**Figure S3.**
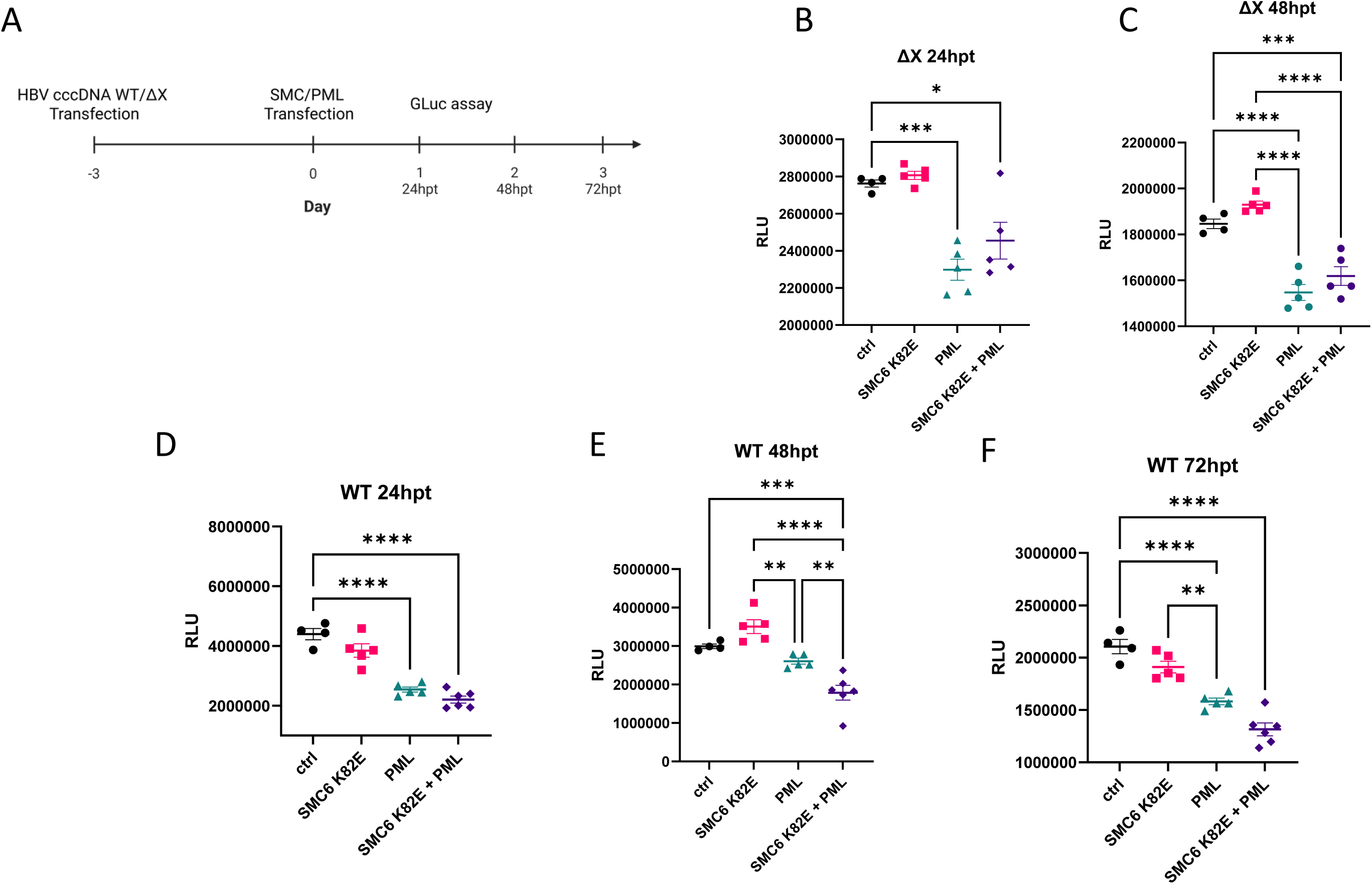
PML-mediated repression of WT HBV cccDNA persists despite dominant-negative SMC6. (A) Experimental timeline. HepG2 cells were transfected with WT or HBx-deficient (ΔX) mcHBV-GLuc cccDNA on day -3 and, on day 0, were transferred to 96-well plates and transfected with control vector, SMC6 K82E, PML-4, or SMC6 K82E plus PML-4. GLuc activity was measured at 24, 48, and 72 hpt. (B-C) ΔX mcHBV-GLuc activity at 24 hpt (B) and 48 hpt (C), showing the early response to SMC6 K82E before the 72-h endpoint shown in Fig. 4. (D-F) WT mcHBV-GLuc activity at 24 hpt (D), 48 hpt (E), and 72 hpt (F). Data are shown as mean ± SEM. Statistical analysis was performed by one-way ANOVA followed by Fisher’s LSD post-hoc test (*P < 0.05; **P < 0.01; ***P < 0.001; ****P < 0.0001; ns, not significant).

**Figure S4.**
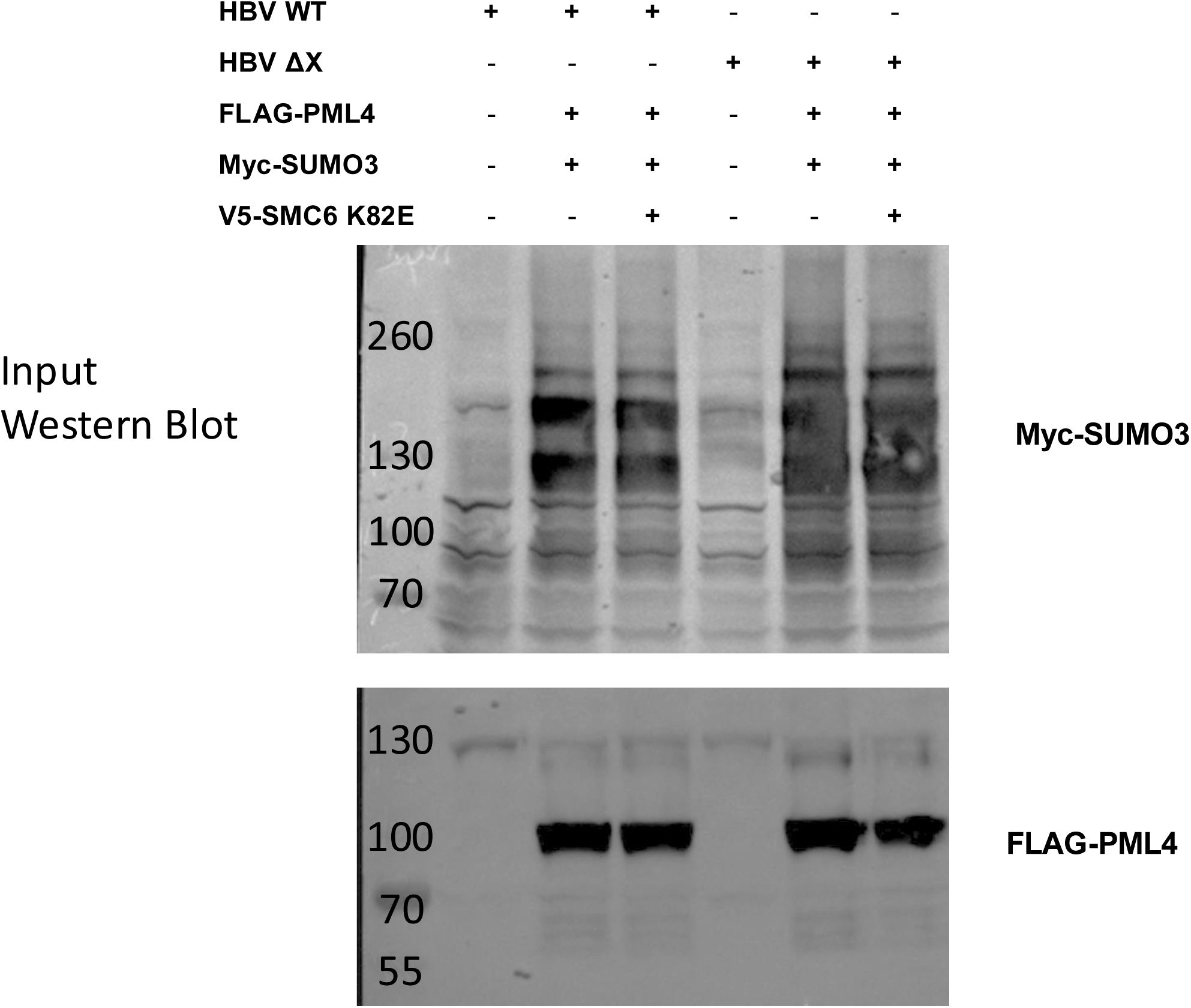
Input controls for the SMC5/6-PML SUMOylation assay. Whole-cell lysates corresponding to the FLAG- immunoprecipitation samples in Fig. 5 were analyzed by immunoblotting to confirm expression of Myc-SUMO3 and FLAG-PML4 across the indicated HBV WT, HBV ΔX, and SMC6 K82E conditions.

**Figure S5.**
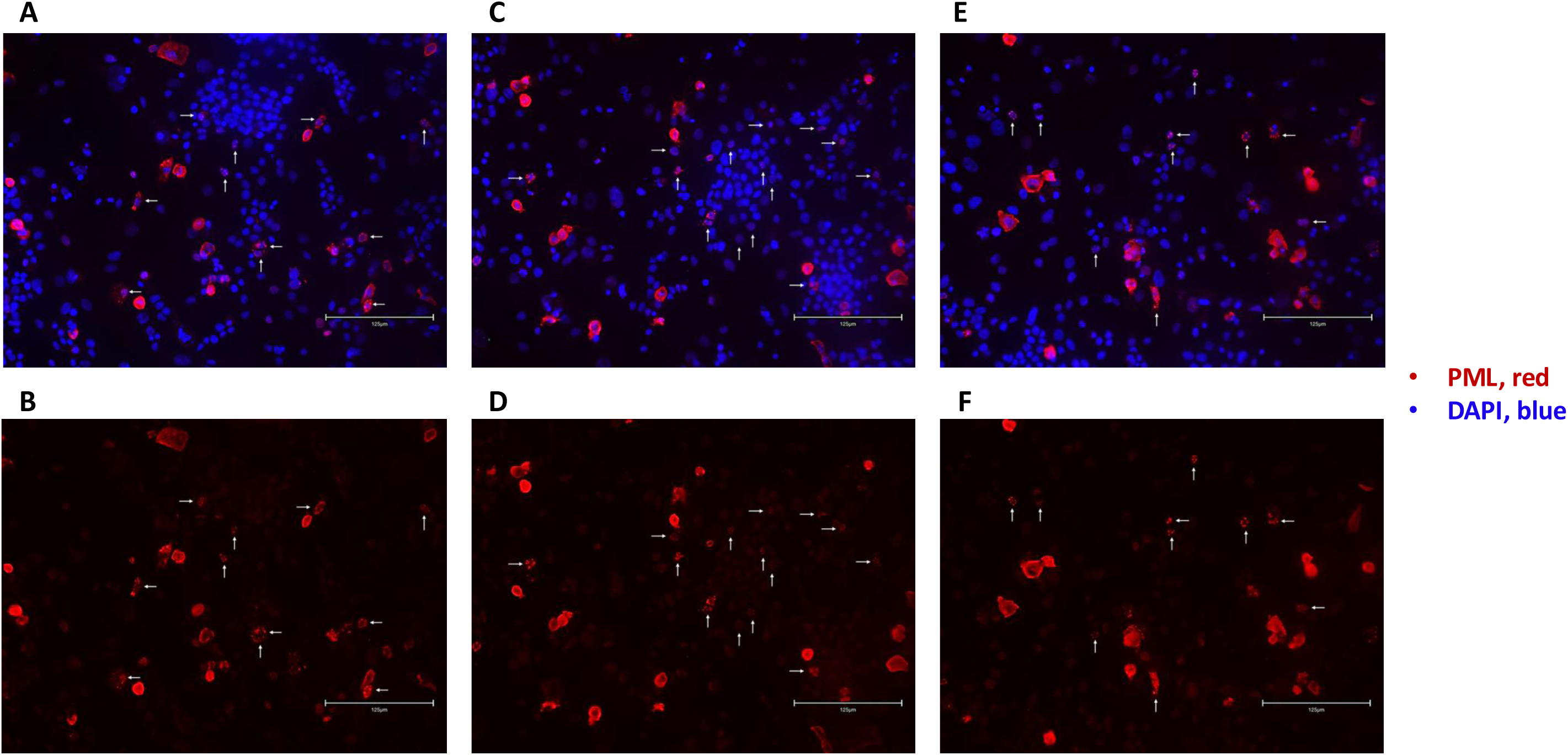
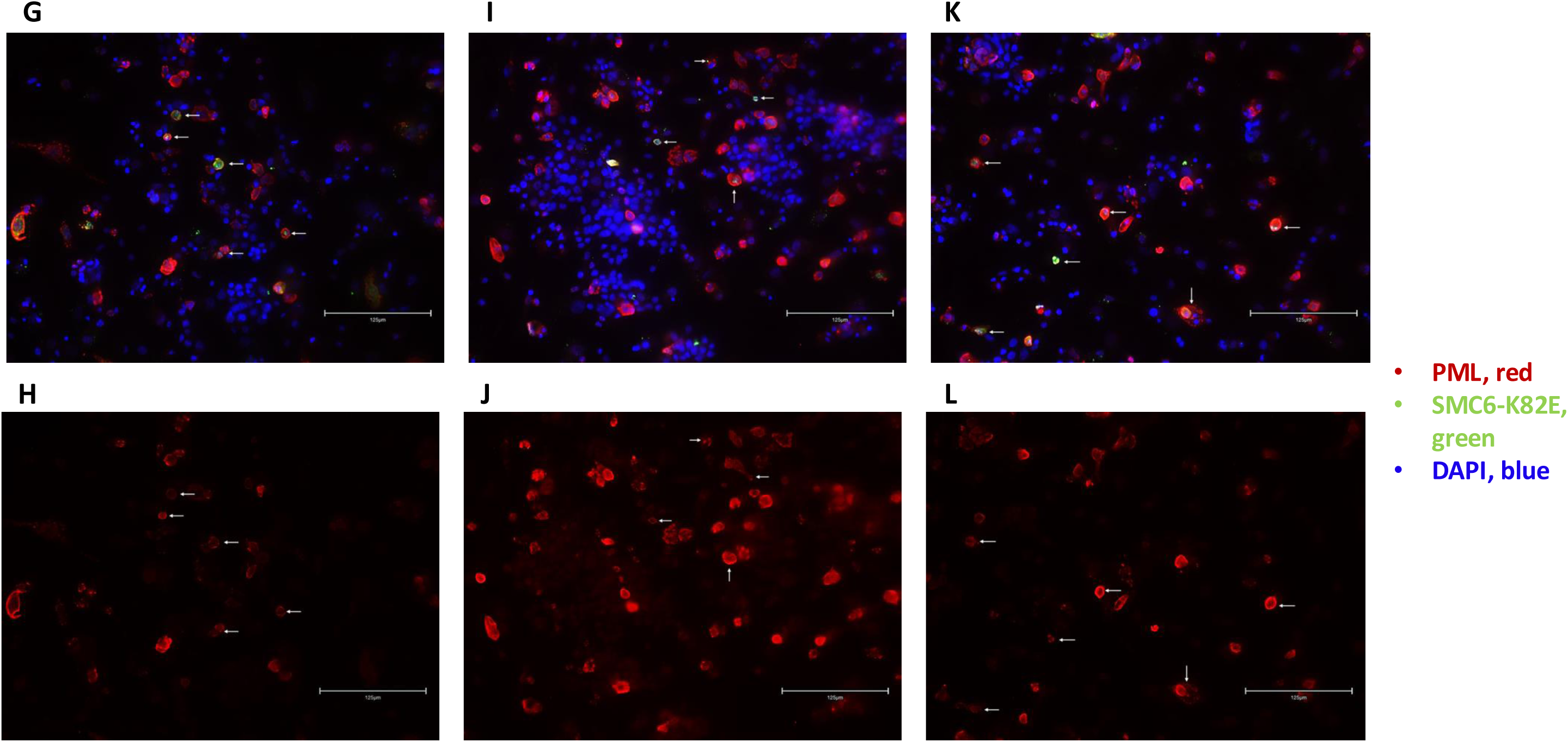
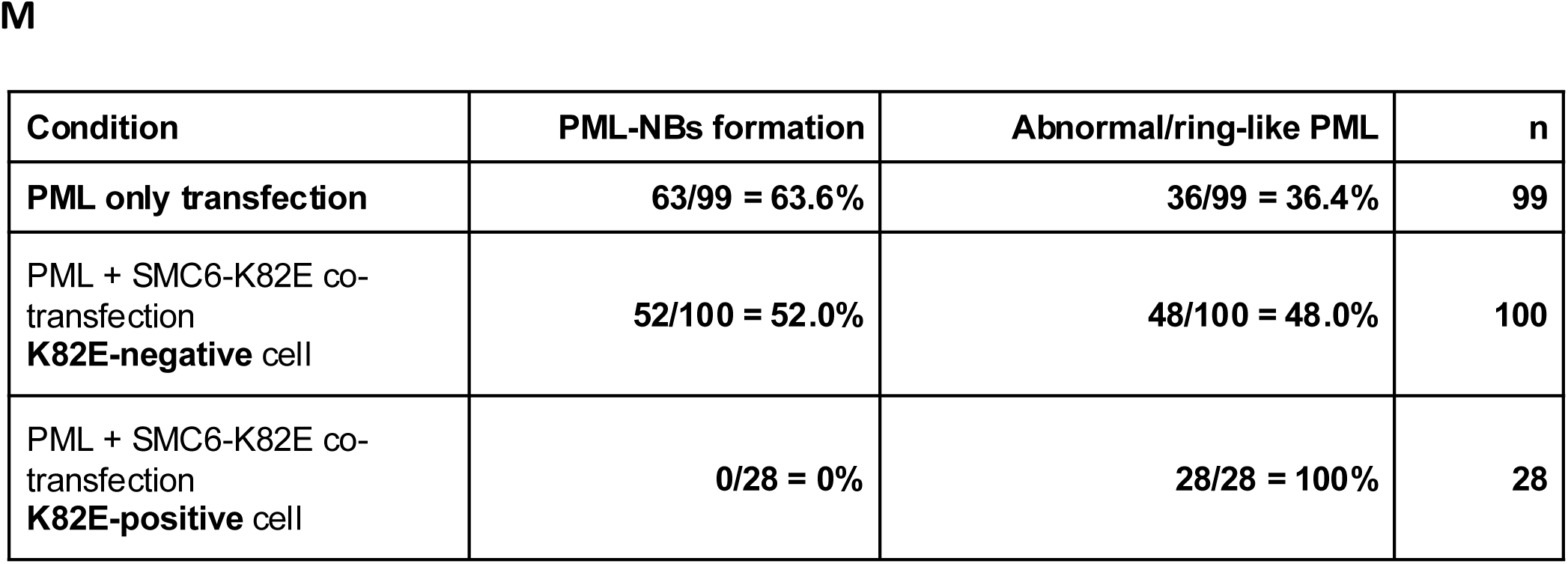
Immunofluorescence fields used for quantification of PML nuclear-body morphology. (A, C, E) Merged PML/DAPI images from three additional fields of HepG2 cells expressing FLAG-PML4 alone; (B, D, F) corresponding PML channels. (G, I, K) merged PML/SMC6-K82E/DAPI images from three additional fields of cells co-transfected with FLAG-PML4 and V5-SMC6 K82E; (H, J, L) corresponding PML channels. PML is shown in red, SMC6 K82E in green, and DAPI in blue. (M) Pooled cell counts across four fields per group: PML-only, 63/99 punctate and 36/99 abnormal/ring-like; PML-positive/SMC6-K82E-negative, 52/100 punctate and 48/100 abnormal/ring-like; PML/SMC6-K82E double-positive, 0/28 punctate and 28/28 abnormal/ring-like. SMC6-K82E positivity was defined by detectable green signal in the same cell. The representative fields in Fig. 6 together with these additional fields comprised the four fields used for the pooled morphology analysis. Arrows indicate representative cells corresponding to the scored phenotype; quantification included all classifiable PML-positive cells within each field. Scale bars, 125 μm.

## Notes

### Competing Interest Statement

The authors have declared no competing interest.

